# A Conditional Cas9 System for Stage-Specific Gene Editing in *P. falciparum*

**DOI:** 10.1101/2025.03.09.642268

**Authors:** Sean T. Windle, Maxwell L. Neal, Fred D. Mast, Stefan H. I. Kappe, John D. Aitchison

**Affiliations:** Department of Global Health, University of Washington, Seattle, WA, USA; Center for Global Infectious Disease Research, Seattle Children’s Research Institute, Seattle, WA, USA; Department of Pediatrics, University of Washington, Seattle, WA, USA

## Abstract

The malaria parasite has a complex lifecycle involving various host cell environments in both human and mosquito hosts. The parasite must tightly regulate gene expression at each stage in order to adapt to its current environment while continuing development. However, it is challenging to study gene function and regulation of essential genes across the parasite’s multi-host lifecycle. Thus, we adapted a recently developed a single-plasmid dimerizable Cre recombinase system for rapamycin-controllable expression of Cas9, allowing for conditional introduction of mutations. We explored rates of gene deletion using varying repair template lengths, showing functionality of donor templates under 250bp for homology-directed repair. As a proof of concept, we conditionally disrupted two uncharacterized genes in blood and gametocyte stages, identifying new stage-specific phenotypes.

**Importance:** As progress towards eliminating malaria has stalled, there is a pressing need for new antimalarials and vaccines. Genes essential to multiple stages of development represent ideal candidates for both antimalarials and vaccines. However, much of the parasite genome remains uncharacterized. Conditional gene perturbation approaches are needed in order to study gene function across the lifecycle. Currently available tools are limited in their ability to perturb genes at the scale required for large screens. We describe a tool that allows for conditional introduction of desired mutations by controlling Cas9 with the DiCre-loxP system. We demonstrate the accessibility of this approach by designing gRNA-donor pairs that can be commercially synthesized. This toolkit provides a scalable system for identifying new drug and vaccine candidates targeting multiple stages of the parasite lifecycle

## Introduction

Malaria is a life-threatening vector-borne disease caused by apicomplexan parasites, with *Plasmodium falciparum* responsible for over 600,000 deaths and 200 million cases globally per year (1). Since the beginning of the COVID-19 pandemic in 2020, progress towards reducing the burden of malaria has halted, with many efforts and interventions being disrupted by the pandemic (2). These setbacks have led cases and mortality to increase from 2019, pushing the goal of eradication even further away. Efforts have been further complicated by the rise of multi-drug resistant strains of *P. falciparum* (3). There is an urgent need for new antimalarials, as well as long-lasting and multistage vaccines. Understanding *Plasmodium* biology across the parasite lifecycle is integral to developing antimalarial treatments, vaccines, and vector-based controls. However, there is still much that is unknown about the underlying biology of the parasite, as roughly one third of the genome remains completely uncharacterized.

The *Plasmodium falciparum* lifecycle consists of multiple stages of development that involve both a mammalian and mosquito host. Beginning with an infectious mosquito bite, parasites migrate to the liver where they infect hepatocytes. Parasites replicate to form tens of thousands of merozoites before egressing out of the host hepatocyte and proceeding to infect red blood cells, initiating the blood stage. Once in the blood cells, parasites develop first into ring-shaped trophozoites, then develop into schizonts, before finally rupturing the red blood cells to release new merozoites, completing a cycle of the blood stage every 48 hours. A subset of these parasites differentiate into committed sexually dimorphic gametocytes, either as a ‘male’ gametocyte or a ‘female’ gametocyte. Upon a subsequent mosquito bite, gametocytes are ingested and rapidly develop into gametes, which fuse to form a zygote in the mosquito midgut before forming oocysts. The parasites then continuously replicate, producing thousands of sporozoites which migrate to the salivary glands of the mosquito, thus completing the cycle. There are multiple bottlenecks within the parasite lifecycle which offer potential opportunities to halt the lifecycle entirely. However, due in large part to the complexity of the Plasmodium lifecycle, the design of screens that allow for the identification of gene candidates for developing new antimalarials and vaccine targets remains highly challenging.

Genome editing in *Plasmodium* offers a means of identifying essential genes and characterizing gene function. With the ability to transfect plasmids, one is able to create specific mutations and knockouts through homologous recombination. Since the introduction of CRISPR/Cas9, Cas9-mediated editing has become the default approach for editing the *Plasmodium* genome. However, *Plasmodium* lacks the non-homologous end-joining DNA (NHEJ) repair pathway, preventing the formation of insertion and deletions (indels) as a result of Cas9 DNA cutting and NHEJ repair (4). Instead, *Plasmodium* parasites almost exclusively repair DNA strand breaks through homology-directed repair (HDR). HDR-mediated editing allows for the introduction of large gene insertions, deletions, or replacements using a gRNA to direct Cas9 cleavage to the target site, and a DNA donor sequence, usually provided by plasmid. Although this approach provides a means of editing the parasite genome with high success, the process can still take many months to generate a single knockout line, as it requires multiple steps of bacterial cloning as well as an extensive period of recovery following transfection. Further, for *Plasmodium* genes essential for blood stage growth, a knockout will be lethal and therefore will not allow for progression onto subsequent stages of development.

Very few genome-wide screens have been performed to identify essential genes in *Plasmodium,* in large part due to the challenges in designing such a system and the limitations inherent in *Plasmodium* biology. A genome-wide *P. berghei* knockout screen using a library of gene targeting vectors was performed, targeting 2,578 genes. In this study, approximately 45% of genes were found to be essential for blood stage growth, while an additional 18% resulted in slowed growth (5). To further elucidate gene essentiality across the lifecycle, 1,300 blood stage viable clones generated in the prior screen were fed to mosquitos to identify gametocyte and mosquito essential genes, and parasites were then allowed to reinfect mice by mosquito bite to identify liver stage essential genes (6). In *P. falciparum,* a PiggyBac transposon screen was done by performing 9,000 individual transfections in order to establish >38,000 unique transposon insertions throughout the genome (7). In this study, approximately 60% of genes were found to be essential to blood stage growth. This was the first genome-wide screen for essentiality in *P. falciparum*, offering numerous insights and further validating the findings of the *P. berghei* screen. However, in neither screens was there the ability to further study the phenotypes of essential genes beyond the blood stage.

A number of approaches have attempted to solve these limitations and simplify the process for characterizing genes across the lifecycle. Rather than performing a gene knockout, genes can be modified to include destabilization domains to control protein levels, as in the ddFKBP-system (8). Alternatively, the RNA can be modified by including a 5’ or 3’ aptamer sequence to be expressed with the gene, allowing for controllable degradation through a TetR-DOZI fusion protein (9, 10). The TetR repressor protein is used for binding to the aptamers, and is controllable through anhydrotetracycline (aTc) in the media, while the DOZI protein is an RNA helicase which diverts RNA to a degradation pathway, thereby repressing translation. The fusion of these allows for strong knockdown of genes tagged with the aptamers when in the absence of aTc.

zzAnother approach for conditional gene disruption utilizes a dimerizable Cre, which consists of two halves fused to FKBP12 and FRB proteins, allowing for dimerization to form a functional Cre recombinase in the presence of rapamycin (11–13). This system allows for chemically induced recombination and deletion of genes flanked by loxP sites. This requires integrating loxP sites on both sides of the target gene and installing silent mutations to disrupt the Cas9 cut site, as well as cloning of the gRNA itself.

These approaches have been used to characterize mechanisms of erythrocyte invasion, developmental stage transition, and drug resistance (14–18). In all of these systems, however, the target gene must be edited to include the respective changes to genomic DNA through homology-directed repair, requiring extensive bacterial cloning and often months of effort to generate a single parasite line. In order to overcome this limitation, we developed a single-plasmid rapamycin-inducible Cas9 system that allows for conditional editing of the *P. falciparum* genome. We first tested our system by reproducing a knockout of the Circumsporozoite Protein (CSP) gene, which is dispensable for the blood stage. To increase the scalability of cloning, the system was adapted to allow for cloning of synthesized gRNA-donor pairs. We then explored the knockout phenotypes of two uncharacterized genes, PF3D7_1454900 and PF3D7_1316300, using this approach, identifying uncharacterized phenotypes in the blood stage and gametocyte stage.

## Results

### Design of a DiCre-inducible Cas9

We first evaluated existing conditional Cas9 systems, including a TetR aptamer-regulated Cas9, and a chemically inducible Cas9 (ciCas9) (9, 10, 19). However, in *Plasmodium*, the TetR aptamer system was too leaky, and the Bcl-xL inhibitor, A115, used for inducing ciCas9 proved too toxic to red blood cells, making them unsuitable for conditional editing. We then explored a dimerizable Cre (DiCre) approach, which has been used for inducible expression of AP2-G to rapidly convert blood stage parasites into gametocytes (18). While we were adapting the DiCre system, another group independently applied a similar strategy for DiCre-mediated expression of dCas9-effector proteins (20). We sought to build on this design to create a DiCre-mediated system for Cas9 activation.

To generate a single-plasmid system for Cas9 editing, we incorporated the loxP-GFP-STOP-loxP cassette including the Hsp86 promoter and terminator into our Cas9 backbone, alongside a constitutively expressed Blasticidin resistance cassette. The GFP sequence in the original loxP-GFP-STOP-loxP cassette was replaced with codon-optimized mNeonGreen (mNG) to improve visibility across all parasite stages. To ensure cleavage occurred only at the specified gRNA target site, we replaced wildtype SpCas9 with a high-fidelity variant, creating the DiCre-Cas9 plasmid (21) (Supp. Figure 1).

In parasites expressing DiCre, transfection with this plasmid followed by rapamycin treatment induces dimerization of the Cre components, generating an active recombinase that excises the STOP cassette, turning off mNG expression and activating Cas9 (Figure 1A). If a gRNA and HDR donor template are present, Cas9 cleaves the target site, and the parasite’s repair machinery uses the donor template to generate the intended mutation, allowing for potential rescue of the edited parasite.

**Figure 1.**
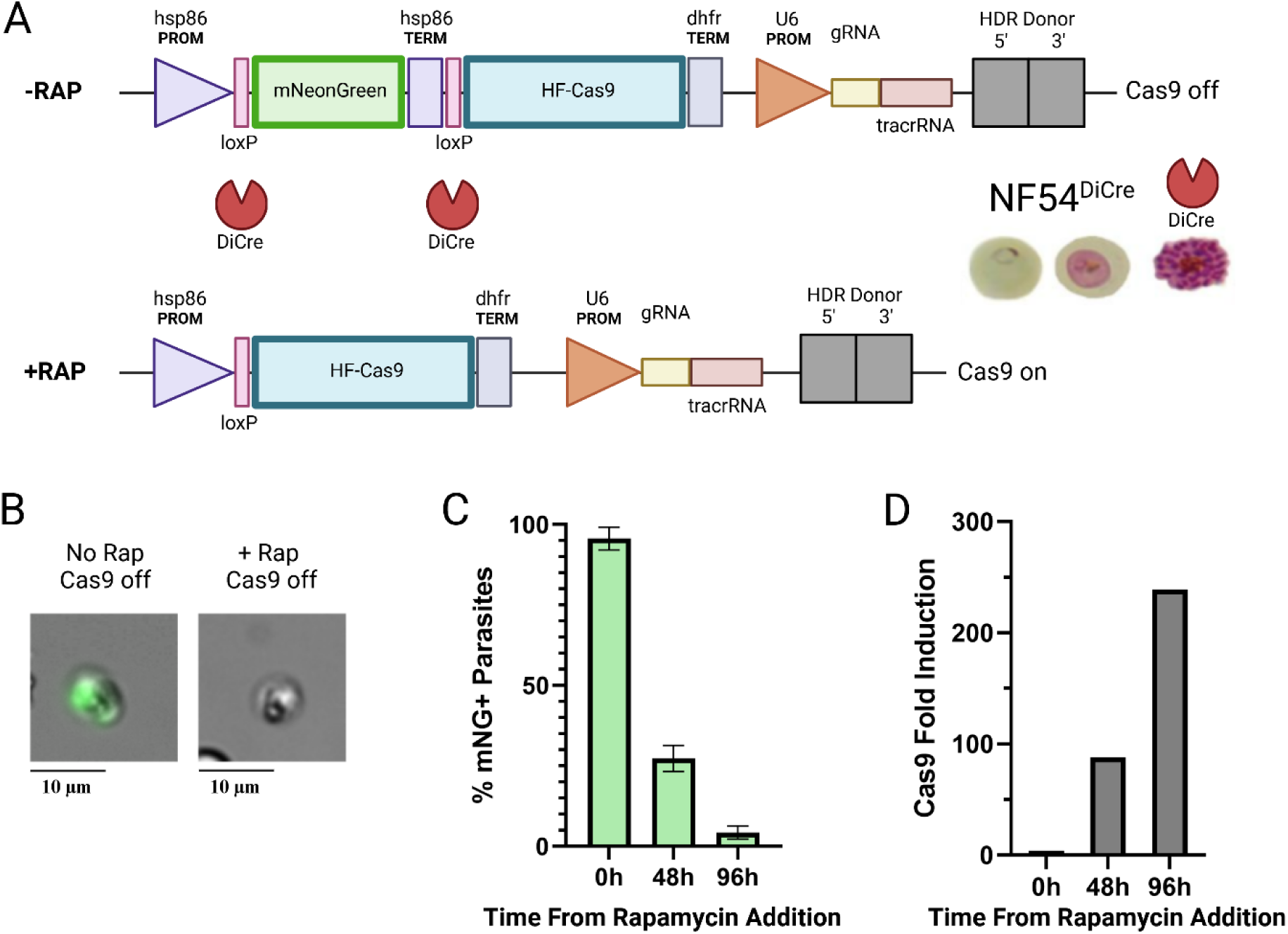
DiCre-inducible Cas9 system. **A)** Schematic showing the DiCre-inducible Cas9 system. The Hsp86 promoter is followed by a loxP-flanked mNeonGreen and Hsp86 terminator, creating a loxP-STOP-loxP cassette upstream of a high-fidelity Cas9. When used in the NF54^DiCre^ parasite line, addition of rapamycin activates DiCre, excising the STOP element between the two loxP sites and activating expression of Cas9. The plasmid also contains a U6 promoter driving gRNA expression, allowing for targeted cleavage by Cas9, and an HDR donor template that allows for repair of the cleaved DNA. **B)** Fluorescent microscopy of representative trophozoites before and after addition of rapamycin, showing that mNG expression has been extinguished. Bar – 10 µm. **C)** Percent of mNG-positive parasites at 0h, 48h, and 96h following addition of 50nM rapamycin. Parasites were tightly synchronized to early ring stage before rapamycin addition. Data shown represents the means of three replicates ± SEM. **D)** Fold-induction of Cas9 relative to uninduced at 0h, 48h, and 96h following addition of 50nM rapamycin. Expression was measured by SYBR Green RT-qPCR assay for Cas9 mRNA at respective timepoints.

Initial testing of this system in an NF54-DiCre parasite line demonstrated highly effective excision of the mNG gene (Figure 1B). Rapamycin treatment at 50nM for 6h maximized excision efficiency without detectable toxicity. As expected, the mNG fluorescence was significantly diminished in most parasites after rapamycin treatment but persisted at a reduced level until the following cycle, likely attributable in part to protein stability following gene excision. By 48h post-treatment, approximately 70% of parasites had lost mNG signal, and by 96h, 96% of parasites showed no mNG signal (Figure 1C). Conversely, Cas9 expression increased 239-fold by 96h post-treatment with rapamycin (Figure 1D). Although rapamycin was removed from the culture at hour 6, it is possible that residual rapamycin may contribute to the continued conversion of parasites from mNG+ to mNG-.

As a proof of concept for inducible Cas9 editing, we incorporated a validated gRNA and donor template targeting the Circumsporozoite Protein (CSP) gene, which is dispensable for both blood and gametocyte stages. This donor contained 550bp homology arms containing the CSP 5’UTR and 3’UTR sequences, but omitted the CSP coding sequence. In this design, addition of rapamycin should extinguish mNG and activate Cas9 expression, leading to Cas9-mediated cleavage at the CSP gRNA target site and subsequent repair through the HDR pathway, with the consequent loss of the CSP coding sequence (Figure 2A). To prevent continuous cleavage following repair, the donor template was designed without the gRNA target sequence. Parasites that fail to repair with the donor should die due to the unrepaired double-strand break.

**Figure 2.**
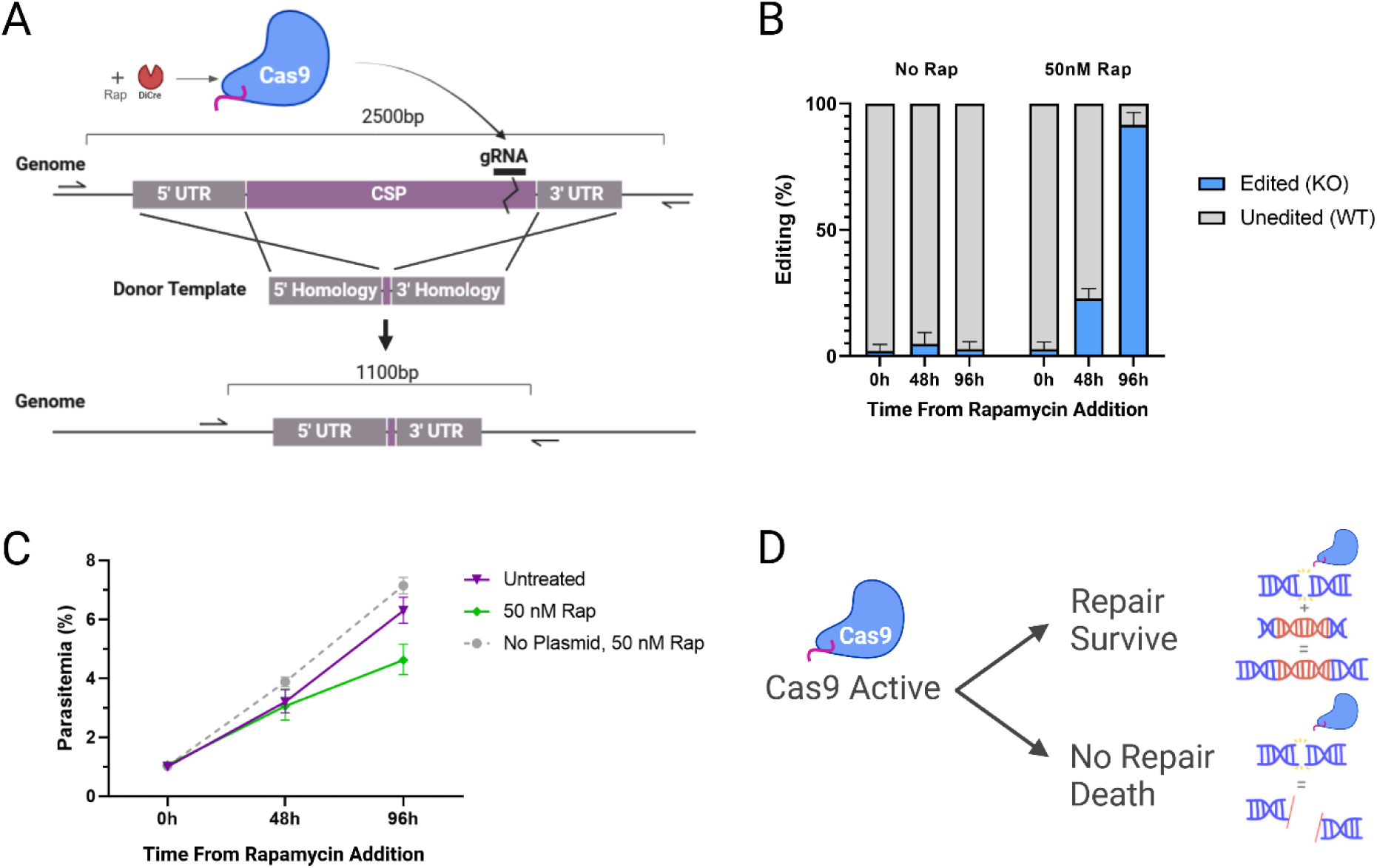
DiCre-inducible CSP editing in blood stage parasites. **A)** Rapamycin activates DiCre, which in turn activates Cas9 expression and gRNA-directed cleavage at the 3’ end of the CSP gene. A donor template with homology to the 5’ and 3’ UTRs of CSP but lacking the coding sequences is used for repair, creating a CSP knockout. Two external primers are used for amplification of the unedited (∼2500 bp) or edited (∼1100 bp) CSP locus, allowing for quantification of editing rates following induction. **B)** Inducible editing rates at 0h, 48h, and 96h after adding 50nM rapamycin or DMSO. Correctly edited alleles contain the intended CSP deletion, while unedited alleles are WT CSP. Data shown represents the means from three experimental replicates ± SEM. (* p < 0.05). **C)** Growth rates following addition of either 50nM rapamycin or 0nM (untreated) to tightly synchronized ring stage parasites at 1% starting parasitemia. Untreated plasmid-free parasites were included as a control. Data shown represent the means from three or more replicates ± SEM. **D)** Diagram showing the binary outcomes following Cas9-induced double strand breaks in the genome. Parasites that repair the break using the donor template survive, while parasites that fail to repair die.

This system was first tested in red blood cells, which were synchronized at the ring-stage of parasite development. Cells were treated with 50nM rapamycin for 6h and the drug was then washed out. Parasites were sequenced at each cycle (48h and 96h) to track the efficiency of editing over time. At hour 0 (no rapamycin), there was typically minimal editing (<5%) from the CSP WT to KO sequence, as measured by PCR and amplicon sequencing. At both 48h and 96h, statistically significant rates of editing were observed, with 22% of parasites containing KO sequences at 48h, and 89% containing KO sequences at 96h (Figure 2B). This temporal delay of editing following rapamycin addition may be in part due to the nature of the inducible system, as well as the activity of parasite repair pathways during development. In this system, DiCre must first be activated, allowing for binding and cleavage of loxP sites, removal of the mNG-terminator, and activation of Cas9 expression. Cas9 must then localize to the gRNA target site and create a double-strand break. Only then can the parasite repair this break using the donor sequence and HDR.

Although no immediate toxicity was observed based on morphology, we were interested in determining whether Cas9 activity or repair had any effect on parasite growth or survival. At 48h, there was no significant difference in parasitemia or morphology between rapamycin-treated and -untreated parasites. However, by 96h, a significant decrease in parasitemia was observed, with 32% lower parasitemia, suggesting Cas9-induced toxicity or a minor growth defect over extended time (Figure 2C). Because double-strand break repair must occur before replication, parasites may not survive or replicate in cases where Cas9 cleavage occurs before or during replication (Figure 2D). This is particularly relevant given that HDR activity is highest during active replication. It is possible that parasites must halt development or replication in order to repair DNA breaks induced by Cas9 during this period, resulting in stalled growth.

We next tested whether inducible editing could occur in gametocytes. Rapamycin was added to committed gametocytes at 3 timepoints (Day 1, Day 5, and Day 7), and editing was then measured on Day 10 (Figure 3A). To eliminate blood-stage parasites, gametocyte cultures were treated with *N*-acetylglucosamine (GlcNAc), which is toxic to late blood stage parasites, for 5 days beginning on day 0 (22). At all three timepoints, gametocytes retained substantial HDR capabilities despite not actively dividing (Figure 3B). By Day 10, up to 82% of gametocytes treated on Day 1 were successfully edited to CSP KO, and up to 55% of those treated on Day 7 were successfully edited, with significant editing observed for all timepoints. Although a small amount of background may originate from residual blood stage parasites, the majority of gametocytes underwent HDR and appeared morphologically normal. These results demonstrate that the DiCre Cas9 system enables inducible editing of gametocytes at moderate to high efficiency.

**Figure 3.**
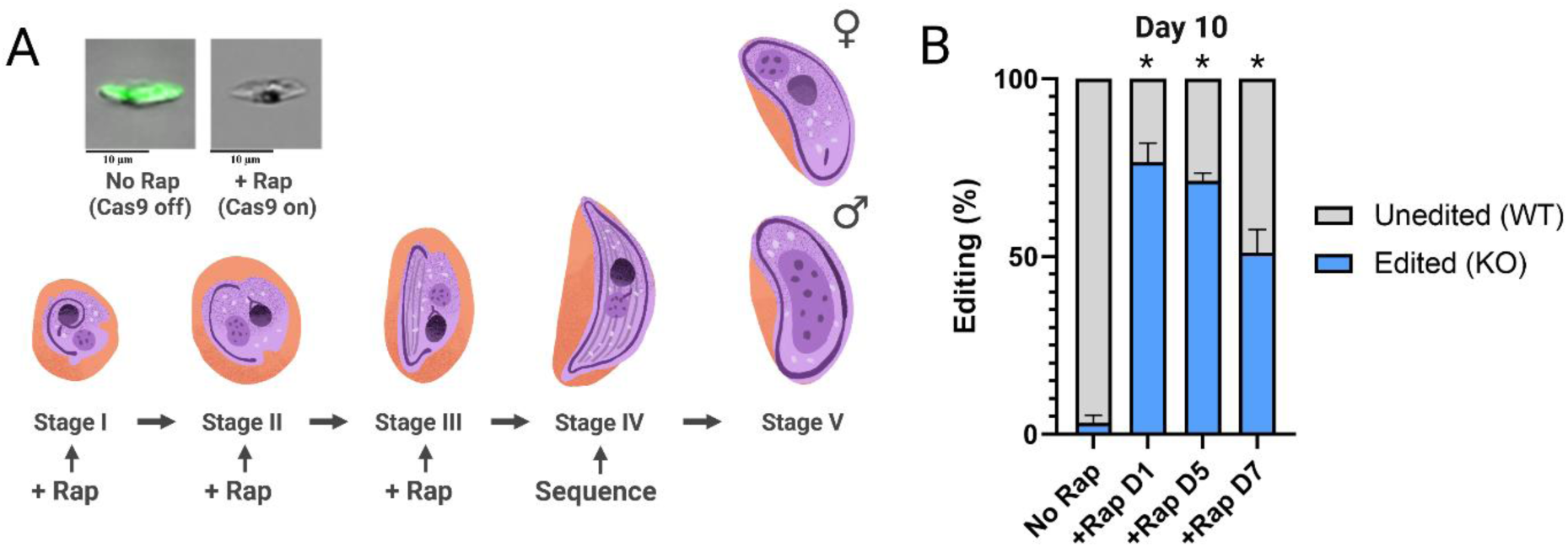
DiCre-inducible CSP editing in gametocytes. **A)** Timeline for inducible Cas9 editing in gametocytes. 50nM rapamycin or DMSO (No Rap) was added on Day 1, Day 5, or Day 7 following gametocyte induction. DNA was isolated from parasites on Day 10 and sequenced. **B)** Efficiency of CSP editing to the KO allele in gametocytes at Day 1, Day 5, or Day 7 after addition of rapamycin or DMSO, as measured on Day 10. Data shown represents the means from three or more replicates ± SEM. (* p < 0.05)

### Improving the Accessibility and Scalability of the Inducible Cas9 System

To improve the scalability of the system, we sought to overcome the limitations associated with cloning individual gRNAs and inserting two fragments encoding the donor sequence’s homology arms, a process that requires multiple rounds of PCRs, ligations, and sequencing for each target site and can take weeks to complete.

Recent CRISPR-based approaches in *S. cerevisiae* and *T. gondii* utilized short (<300bp) gene fragments to perform HDR screens (23). This strategy allows one to synthesize the 20bp gRNA as well as its respective HDR donor, pairing the synthesis and cloning steps. However, in these studies, very short (30-75bp) homology arms were required in order to fit into the <300bp fragments. Advances in DNA synthesis now allow for the affordable synthesis of 500–1000 bp DNA fragments, overcoming this limitation. To adapt this approach, we modified the DiCre-inducible Cas9 plasmid by removing the 85 bp tracrRNA originally following the gRNA sequence (Figure 4A). This modification was necessary for contiguous pairing the gRNA and donor sequences without intervening sequences between the U6 promoter, the gRNA, and the tracrRNA. The tracrRNA can then either be included with the gRNA-donor synthesis, or cloned separately using Type IIS restriction enzymes. Given the affordability of DNA synthesis, we opted to include the tracrRNA in all synthesized gRNA-donor pairs. An additional benefit of this approach is the potential for generating gRNA-donor libraries targeting tens or hundreds of genes simultaneously, providing a scalable and efficient alternative for high-throughput genome editing. We assessed the efficiency of HDR-mediated gene deletion using the same gRNA and CSP donor, but varying homology arm lengths (∼80bp, ∼200bp, or ∼500bp for each arm) (Figure 4B). At 96h, all 3 donor sizes had significant increases in editing efficiency with knockout rates at 15%, 49%, and 80% respectively. Notably, the 80bp homology arms resulted in significantly lower editing efficiency compared to 200bp or 500bp. It is typical that reduced homology arm length results in reduced HDR in other systems, and in this case, this reduced efficiency may also be compounded by a limited accessibility of the episomal repair template for HDR, particularly when homology arms are short (24–26).

**Figure 4.**
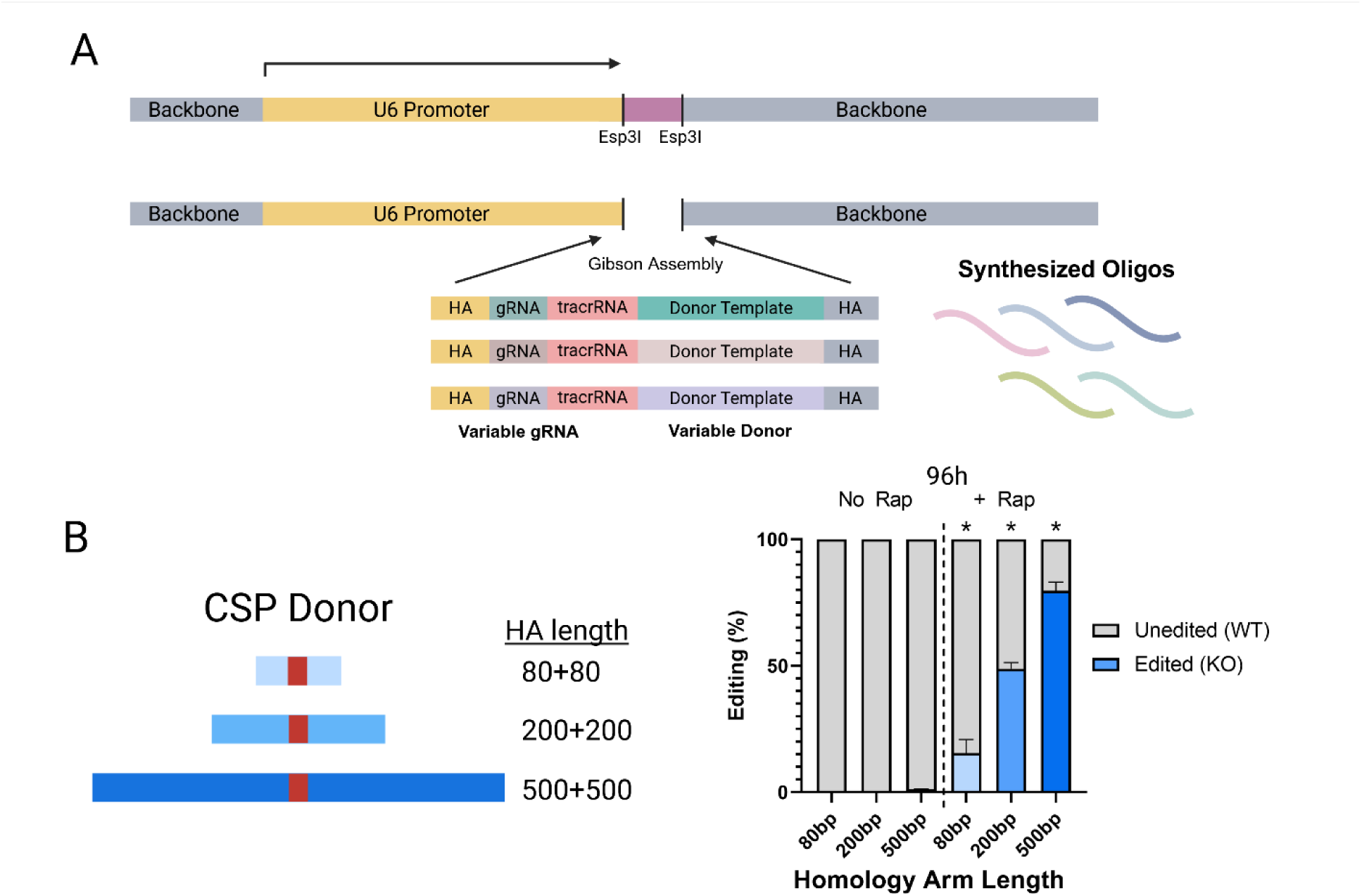
Adaptations to the DiCre-inducible Cas9 system for scalable approaches. **A)** The DiCre-Cas9 plasmid has had the tracrRNA removed, allowing for insertion of a complete gRNA-tracrRNA-donor template oligo by Gibson assembly. Each synthesized oligo contains 25-30bp homology arms (5’ and 3’) for cloning by Gibson assembly, a specific 20bp gRNA, an 82bp universal tracrRNA, and a variable-length donor template. gRNA-tracrRNA-donor oligos can be synthesized individually, in oligo pools, or in arrays. **B)** Testing CSP editing efficiencies using donor homology arms of varying lengths. Three donor templates with homology arm lengths of 80bp, 200bp, or 500bp were used to delete the open reading frame of CSP. Tightly synchronized rings were treated with 50nM rapamycin or DMSO and sequenced 96h later. Data shown represent the means from three replicates ± SEM. (* p < 0.05)

Although donor DNA synthesis is an affordable choice, the highly AT-rich nature of *P. falciparum*’s genome presents a challenge in synthesis, often restricting homology arm lengths to ∼100bp. Despite this constraint, commercial synthesis streamlines molecular cloning, and facilitates the development of pooled or arrayed screens. Altogether, these results demonstrate that DiCre-inducible Cas9 is a robust and scalable system that can be used for efficient editing even with small donor templates.

### Measuring the Phenotype of Gene Candidates in Blood and Gametocyte Stages using DiCre-inducible Cas9

Using transcriptomics datasets and a gene regulatory network model we developed, we identified two previously uncharacterized genes, PF3D7_1454900 and PF3D7_1316300, that are highly expressed in gametocytes with expression higher in female gametocytes (Figure 5A) (27). Both genes were putatively found to be putatively indispensable in a previous PiggyBac transposon screen (7). PF3D7_1454900 encodes a 154 amino acid residue protein, while PF3D7_1316300 encodes a 104 amino acid residue protein. Both genes are moderately conserved among *Plasmodium* species, with at least 50% sequence similarity between their *P. berghei* and *P. yoelii* orthologs.

**Figure 5.**
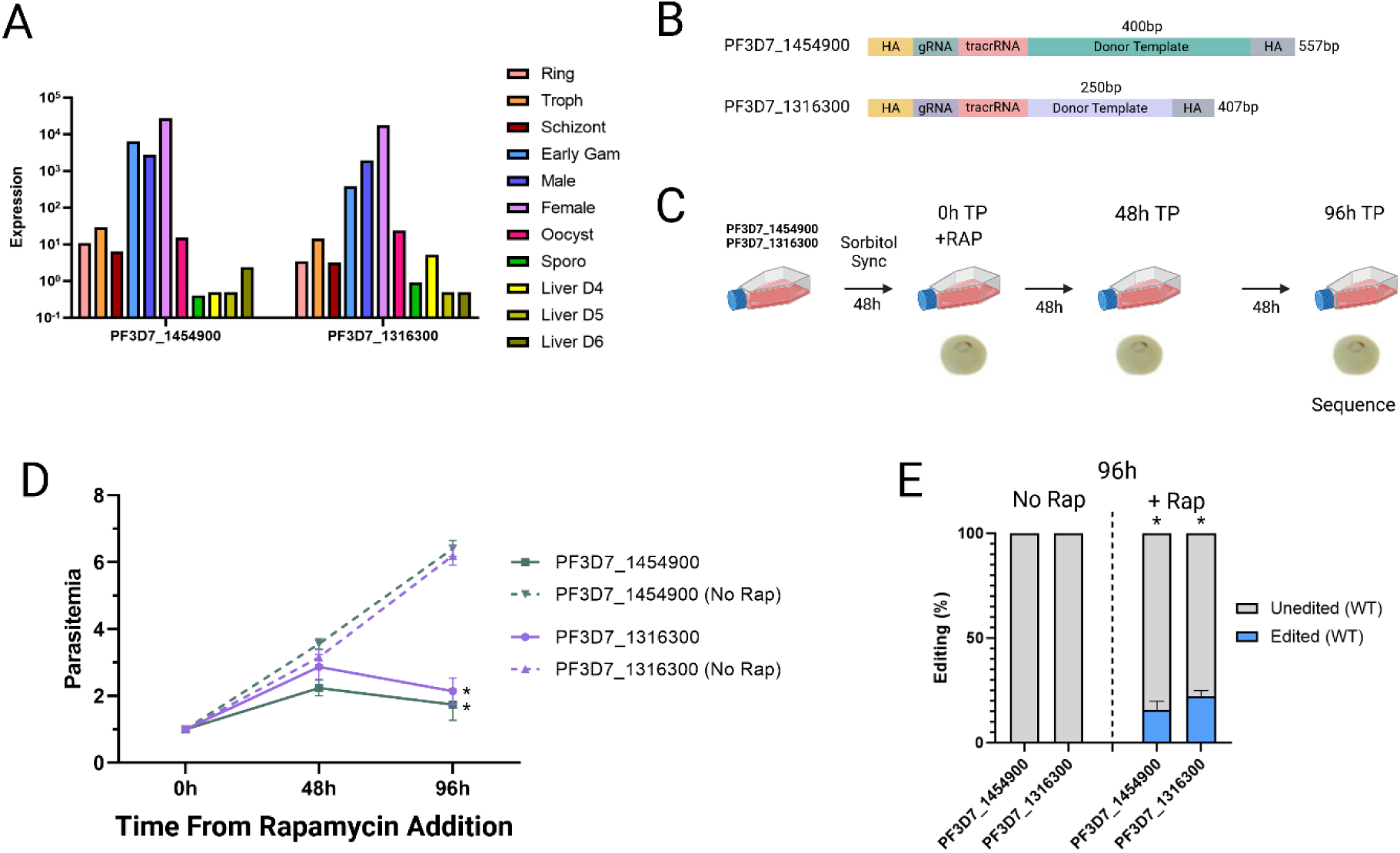
Measuring blood stage essentiality of two gene candidates using DiCre-inducible Cas9. **A)** Normalized gene expression over the parasite lifecycle from compiled transcriptomic datasets for PF3D7_1454900 and PF3D7_1316300. We used 11 broad timepoints, with 3 timepoints in the blood stage, 3 timepoints in the gametocyte stage, 2 timepoints in mosquitos, and 3 timepoints in the liver stage. PF3D7_1454900 is putatively essential by piggyBac screen, whereas PF3D7_1316300 is putatively dispensable by piggyBac screen (7). **B)** Synthesized gRNA-donor pairs for gene candidates PF3D7_1454900 and PF3D7_1316300. The synthesized oligo for PF3D7_1454900 used a 400bp donor template (200bp for 5’, 200bp for 3’), with a total length of 557bp. The synthesized oligo for PF3D7_1316300 used a 250bp donor template (141bp for 5’, 109bp for 3’), with a total length of 407bp. **C)** Timeline for testing blood stage phenotypes. Each transfection was first tightly synchronized to early ring stage parasites before treatment with 50nM rapamycin. Cultures were maintained for 96h (2 cycles), at which time parasitemia was assessed and sequencing was performed to quantify successful editing. **D)** Parasitemia following rapamycin-induced targeting of PF3D7_1454900 and PF3D7_1316300 at 96h post treatment, with and without rapamycin. Data shown represent the means from three replicates ± SEM. (* p < 0.05) **E)** Efficiency of PF3D7_1454900 and PF3D7_1316300 editing to generate the KO alleles in blood stage parasites 96h post rapamycin treatment, or no rapamycin treatment. Data shown represent the means from three replicates ± SEM. (* p < 0.05)

Taking advantage of the accessibility of paired gRNA-donor synthesis, we designed gRNAs paired with the longest HDR donor possible for DNA synthesis (400bp for PF3D7_1454900, and 250bp for PF3D7_1316300) (Figure 5B). Rather than whole gene deletion, we engineered donor sequences to generate a small 8bp deletion and frameshift mutation, while simultaneously disrupting the gRNA site. To assess editing efficiency in blood stage parasites, we induced Cas9 with rapamycin in 0-3h synchronized ring stage parasites and measured parasitemia and editing efficiencies at 96h (Figure 5C). If the frameshift is not deleterious, even low efficiency editing should eventually result in a largely pure population of edited parasites. By 96h post-rapamycin, parasites appeared morphologically normal for both PF3D7_1454900 and PF3D7_1316300. However, in both cases, the parasitemia was significantly reduced, with decreases of 73% and 65%, respectively (Figure 5D). As observed with CSP, parasites continued to grow normally until the following cycle, at which point they either arrested in growth or stalled in development. The proportion of edited parasites in both PF3D7_1454900 and PF3D7_1316300 was 15.7% and 22.3% respectively, suggesting some amount of negative selection may be acting on the mutants (Figure 5E). Considering that both conditional knockouts appeared morphologically unaffected, there may be some impairment in the ability for merozoites to invade, leading to a more sizeable decrease in parasitemia.

To assess gametocyte phenotypes, gametocyte commitment was induced by overgrowth, and a pure gametocyte population was selected for using 50mM GlcNAc. Parasites were treated with rapamycin at Day 5 following removal of GlcNAc and analyzed at Day 14, prior to peak expression (Figure 6A). For both genes, rapamycin treatment significantly reduced the number of viable gametocytes (Figure 6B). While some loss may be due to gRNA inefficiencies, a higher proportion of surviving gametocytes would be expected if this were the primary factor. Morphologically, many parasites resembled dying gametocytes in late stages (Stages 4 to 5) of gametocyte development, with the most pronounced effect in female gametocytes for both genes. These findings suggest essentiality of PF3D7_1316300 and PF3D7_1454900 in mid-to-late gametocytes, particularly in female gametocytes.

**Figure 6.**
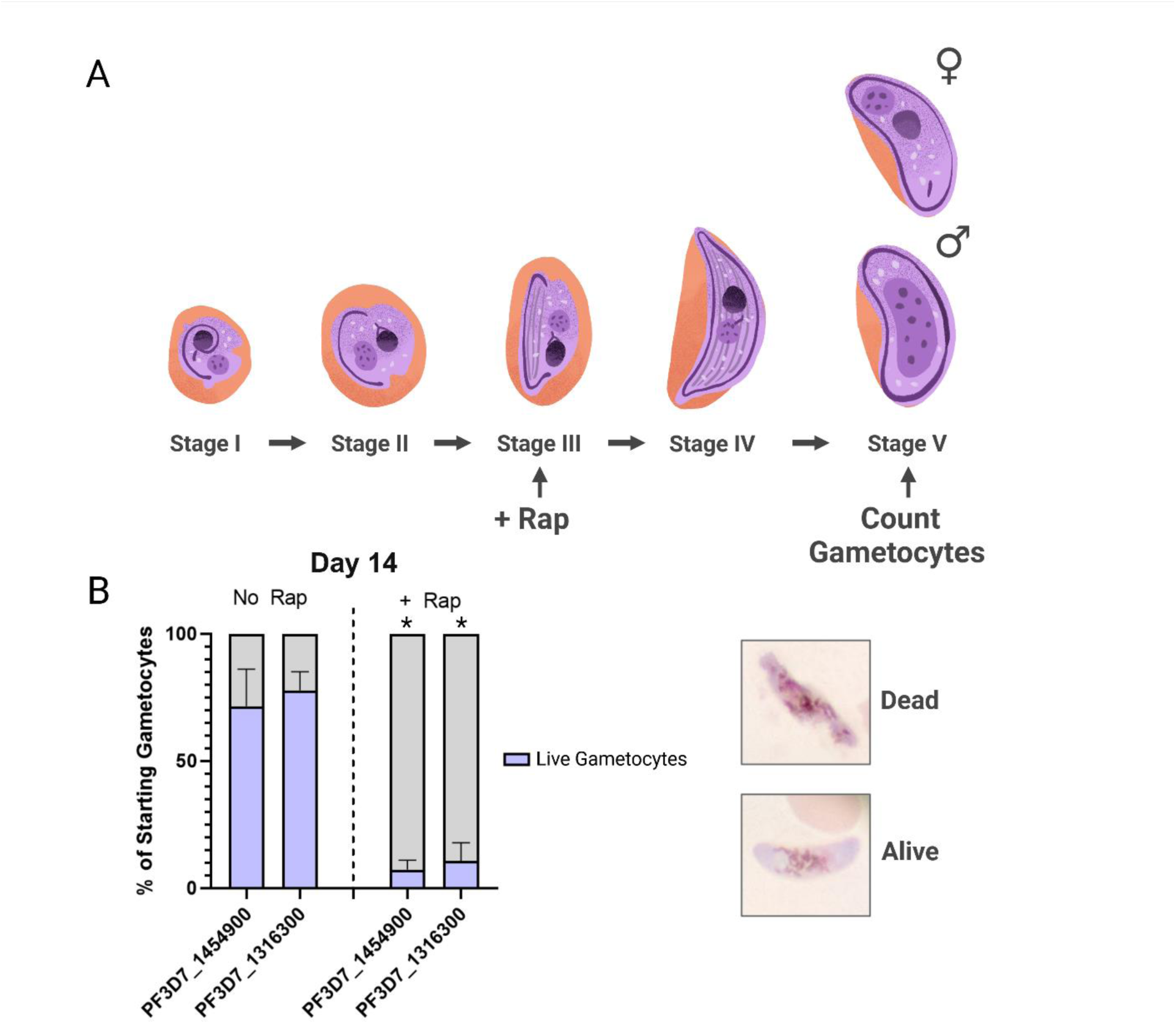
Measuring gametocyte-stage phenotypes in parasites with KOs of two gene candidates using DiCre-inducible Cas9. **A)** Timeline for rapamycin treatment and sequencing to measure gametocyte phenotypes. Induced gametocytes were treated with 50mM GlcNAc for 5 days. On Day 5, 50nM of rapamycin was added and gametocytes were cultured as normal until Day 14, at which point gametocytes were counted by Giemsa staining. **B)** Proportion of starting (Day 5) gametocytes with or without rapamycin induction, counted on Day 14. Live gametocytes retained normal morphology, whereas dead gametocytes began to break apart, condense, or display abnormal morphologies. Data shown represent the means from three replicates ± SEM. (* p < 0.05)

## Discussion

The development of conditional systems for genetic manipulation in *P. falciparum* has expanded functional gene characterization across its lifecycle. We sought to improve upon these strategies by developing a conditional Cas9 system that enables introduction of any desired mutation in an accessible and scalable manner. Rather than requiring multiple independent cloning steps, both the gRNA and its respective donor can be commercially synthesized and cloned in a single step. Using the absence of mNeonGreen signal as an indirect marker of Cas9 expression, parasite phenotypes can be measured by fluorescence microscopy or flow cytometry, as well as sorted by FACS. Our system enables inducible editing in both blood and gametocyte stage parasites, can be applied using small homology arms for Cas9 donor templates, and can be readily scaled up using a paired gRNA-donor approach. We confirmed that the DiCre-Cas9 system facilitates conditional knockout of genes and provides measurable phenotypes in both blood and gametocyte stages. Furthermore, we identified stage-specific essentiality phenotypes in two previously uncharacterized *P. falciparum* genes, PF3D7_1316300 and PF3D7_1454900.

Both conditional knockouts exhibited modest blood stage phenotypes, but had stronger defects in gametocyte stages. These results are in agreement with transcriptomic data, showing peak expression in mid to late gametocytes, particularly in female gametocytes, where we observed the most severe phenotype. Notably, a recent study identified the *P. yoelii* ortholog of PF3D7_1454900 (PY17X_1322400) as a putative apical polar ring (APR) partner protein using TurboID labeling of APR2 and ARA1 (28). These proteins localize to the apical polar ring in *P. yoelii* ookinetes, suggesting a role in microtubule structure and regulation. PY17X_1322400 knockout parasites displayed decreased ookinete motility and near complete deficiency in oocyst development in the mosquito midgut. While the genes appeared dispensable in blood and gametocyte stages in *P. yoelii*, this may be due to inherent differences in parasite biology between *P. falciparum* and *P. yoelii*. For instance, gametocytes in *P. falciparum* require an additional 7 to 9 days to mature, and undergo more distinct morphological development than *P. yoelii*. Additionally, *P. falciparum* gametocytes form a complete inner membrane complex with elongated microtubules whereas *P. yoelii* forms incomplete inner membrane complexes during gametocyte development (29, 30). Based on the available evidence, PF3D7_1454900 may be involved in microtubule formation during brief periods of merozoite invasion and throughout gametocyte development, consistent with the observed phenotypes. However, more thorough study of both PF3D7_1316300 and PF3D7_1454900, is needed to elucidate their precise functions.

This study also evaluated the efficiency of different homology arm lengths for homology-directed repair and editing. CSP editing was least efficient with 80 bp homology arms and most efficient with 500 bp arms, consistent with findings from *P. berghei* and mammalian Cas9 editing systems (25, 26). Although our donor templates are provided as episomal plasmids, they likely form concatemeric structures with unknown copy numbers (31). While *P. falciparum* lacks non-homologous end joining pathways and preferentially repairs double-strand breaks via HDR, its efficiency remains lower than that of *S. cerevisiae*, where near-complete editing can be achieved with just 50 bp homology arms (24, 32). A limitation of our system is the inability to completely distinguish poor repair efficiency from deleterious phenotypes when using donor templates with shorter homology arms, an issue also observed in *S. cerevisiae* (33). This can be partially addressed by examining additional timepoints and continuing rapamycin treatment, while measuring parasitemia and active growth. Editing efficiency may also be improved by implementing a dual-gRNA strategy, allowing for the complete deletion of the target sequence and subsequent repair by HDR (i.e., 5’ and 3’ ends) (34, 35).

Currently, our DiCre Cas9 system is limited to parasite lines already expressing DiCre, which has only been established in NF54 / 3D7 lines (13, 36, 37). Due to the large size of the DiCre cassette, incorporating it into our plasmid would likely be impractical. However, sequential transfections – first introducing a plasmid carrying the DiCre cassette, followed by the DiCre Cas9 plasmid, with dual selection – could potentially increase its applicability.

There is great value in a tool capable of conditionally editing the parasite genome across the entire lifecycle. However, extending this system to the mosquito and liver stages would require genomic integration to ensure efficient passage through the mosquito vector. One potential approach is to use Bxb1 integrase with attP/attB sites to stably integrate the plasmid, including the gRNA and donor template. However, the only established DiCre-attB parasite line at this time is deficient in gametocyte development, prohibiting study of any stages outside of the blood stage (38, 39).

A recently published approach introduced a barcoded loxP intron into the start of an open reading frame to generate a frameshift mutation, disrupting the open reading frame upon DiCre induction (40). This system also utilized commercial DNA synthesis, allowing for scalable genetic screens coupled to barcode sequencing. However, this system is inherently limited to frameshift knockouts and still requires generating the initial parasite lines before conducting experiments. Additionally, the introduction of the loxP-intron-loxP may alter function or be deleterious in certain genes. Nevertheless, this method provides a complementary approach that allows for scalable phenotyping of essential genes across the parasite lifecycle.

Our system represents the first the inducible Cas9-HDR genome editing tool for *P. falciparum*. Beyond gene deletions, this system is also amenable to precise genome modifications, including the introduction of SNPs, for functional analysis of resistance mutations. This system could also be adapted for genetic screens, either in a pooled or an arrayed format, to edit multiple genes or sites simultaneously.

## Methods

### Plasmid construction

The DiCre-Cas9 plasmid was assembled by cloning using Gibson assembly, combining the BSD cassette and loxP-GFP-loxP-dCas9 cassette with the backbone containing a U6 promoter and ampicillin resistance cassette. The BSD and loxP-GFP-loxP-dCas9 cassettes were amplified from the DiCre-dCas9-effector plasmids, kindly gifted by Jun Miao (20). The U6-amp backbone was amplified from the Linker3 plasmid, kindly gifted by Ashley Vaughan (41). Both amplicons contained at least 20-bp of homology to each other. dCas9 was then replaced with a high-fidelity Cas9 using Gibson assembly. The HF-Cas9 was amplified from the VP12 plasmid, kindly gifted by Keith Joung (21). A codon-optimized mNeonGreen was amplified from a plasmid kindly gifted by Ashley Vaughan. GFP was replaced with the mNeonGreen by Gibson assembly, using at least 20-bp of homology to each other.

For gRNA cloning of single guides, two complementary oligos were designed, with a 5’ TATT overhang and a 3’ CTGC overhang, allowing for insertion into Esp3I-digested plasmids. Oligos were first annealed by mixing 1uL of 10x NEB T4 ligase buffer and 1uL 100uM of each primer in a 10uL reaction using following conditions: 95C for 5 minutes, gradual decrease to 25C at a rate of 0.1C per second. Annealed oligos were then diluted 1:200 with water. At least 50 ng of Esp3I-digested plasmid was used with 1uL of diluted annealed oligos, mixed with 0.5uL of NEB Quick Ligase, 5uL of 2x Quick Ligation mix, remainder water for a 10uL reaction. The reaction was left at room temperature for 5-30 minutes before transformation. 2uL of ligation mix was used for transformations.

For donor sequence cloning, the 5’ HDR homology arms and 3’ HDR homology arms were amplified from NF54 genomic DNA, with overhangs for Gibson assembly. Plasmids were either digested by a single-cutter enzyme or amplified with respective overhangs for Gibson assembly. The three components were mixed with 2x Gibson assembly Master mix or 2x NEBuilder Master mix with a ratio of 1 vector: 5 insert 1: 5 insert 2, using a minimum of 50ng of linear vector. 1-3uL of mix was used for transformations.

For combined gRNA-donor cloning, the paired gRNA and donor were synthesized by commercial DNA synthesis (Twist Biosciences or GenScript), containing at least 20 basepairs of Gibson homology arms, the gRNA, tracrRNA, and respective donor sequence. Sequences can be found in Supplemental Text 1-4. An Esp3I-digested DCL3 plasmid with the tracrRNA removed was used for Gibson assembly, with a minimum of 50ng of linear vector and 5x of the synthesized oligo. 1-3uL of mix was used for transformations.

A complete list of primers can be found in Supplementary Table 1.

### Parasite culture

*P. falciparum* parasites were maintained in complete medium consisting of RPMI-1640, 25mM HEPES, 2.4mM sodium bicarbonate, 1% Albumax II, and 2% gentamicin, as previously described (42). Parasites were kept at 37C in a 90% N, 5% CO2, 5% O2 environment. Hematocrit was maintained between 4-5% using washed O+ human donor red blood cells. Synchronizations were performed using 5% sorbitol to achieve 0-3h synchronization, as previously described (43). Parasitemia and morphology were checked by Giemsa stain. For routine culture, an NF54 parasite line that was passed through the complete parasite lifecycle was used to ensure viability at each stage of development. For all DiCre experiments, an NF54-DiCre line capable of mosquito transmission was used (13). The NF54-DiCre line was kindly gifted by Moritz Treeck.

Gametocytes were induced starting with tightly synchronized ring stage parasites at 1% parasitemia at 5% hematocrit. Cultures were kept at 37C in a 90% N, 5% CO2, 5% O2 environment, using a slide warmer during media changes. Media consisted of RPMI-1640, 25mM HEPES, 2.4mM sodium bicarbonate, and 10% human sera. Media was changed daily for 14 days, adding pre-warmed media and ensuring not to disrupt the settled cells. For pure gametocyte cultures, 50mM of GlcNAc was added to culture for 5 days before being removed.

For induction of Cas9, 50nM of rapamycin was added to cultures before being removed 6h later and replaced with complete media.

### Transfection

Transfections were performed by ring-stage electroporation using a Bio-Rad Gene-Pulser II Electroporator set to 0.31kV and 950 μF capacitance, as previously described (44). 50-100ug of total plasmid DNA was used per transfection, mixed with cytomix (120 mM KCl, 0.15 mM CaCl2, 2mM EGTA, 5 mM MgCl2, 10 mM K2HPO4/KH2PO4 25 mM HEPES, pH 7.6). Parasites were synchronized by sorbitol to attain approximately 5% parasitemia at ring-stage. The following day, parasites were pelleted, washed with cytomix, and mixed 1:1 with the plasmid-cytomix solution. Parasites were transferred to a 0.2cm electroporation cuvette before being electroporated. Electroporated parasites were immediately moved to pre-warmed flasks with 6mL CM and 125uL fresh packed RBCs. The flask was then gassed with 90N/5O/5CO2 mixture before being placed in an incubator.

Media was changed for two days (one complete parasite cycle) before drug was added. For original L3 plasmids, 2.5 nM WR99210 was used and maintained in media for 5 days. For all other plasmid transfections, 2.5 μg/mL Blasticidin was used and maintained in media for 6 days. Parasites were maintained with media changes every other day after until visible by Giemsa smear. For DCL3 transfections, if no mNeonGreen signal could be detected using fluorescence microscopy after parasite recovery, Blasticidin was returned to media until fluorescent parasites were pure.

### DNA Extractions

Genomic DNA from parasites was isolated using GeneJET Whole Blood Genomic DNA Purification Kits (Thermo Fisher). Prior to isolating, a minimum of 250uL packed infected red blood cells were spun down and resuspended with 1x PBS + 0.015% saponin, set at room temperature for 5-10 minutes, and then pelleted. The supernatant was discarded and the remaining parasite pellet was used following the manufacturer’s protocol. For the final elution, 50uL of warm (37-50C) water was used in place the provided elution buffer, incubating for 5 minutes at room temperature before spinning at 13,000 x g for 1.5 minutes. The purified DNA was then quantified by Nanodrop.

### DNA Sequencing

Plasmids and transfected parasites were amplified using PrimeStar or PrimeStar Max polymerase. To measure genomic editing, 50ng of isolated genomic DNA and primers external to the repaired DNA were used. PCR products were purified by column purification or gel extraction followed by column purification.

All sequencing was done by Plasmidsaurus. For plasmid cloning, whole plasmid sequencing was used. For PCR sequencing, linear amplicon sequencing was used, allowing for quantification of the editing and non-edited DNA from the raw reads. Edits were quantified by by counting the total number of occurrences of the edited and unedited DNA in raw reads.

### RT-qPCR

RT-qPCR for quantification of Cas9 expression was performed using the Sigma-Aldrich One-Step SYBR Green Quantitative RT-qPCR Kit. 100pg of genomic DNA was used. Two validated primers internal to Cas9 were used, with final concentrations of 200 nM, following the manufacturer’s protocol.

### Fluorescent Microscopy

Infected red blood cells were stained in PBS with 10 μM Hoechst 33342 and incubated at 37C for 10 minutes before washing twice and being resuspended in complete media. Cells were then moved to a slide and a coverslip was added. The cells were viewed using a Nikon Eclipse fluorescent microscope at 400-1000x total magnification. For Hoechst detection, the UV filter was used and for GFP or mNeonGreen detection, the blue filter was used. Parasitemias were quantified from the presence or absence of Hoechst and mNeonGreen, depending on the application.

## Acknowledgements

We thank Lucia Pazzagli for her thoughtful discussion and feedback, and Chloe Chan for her artistic contributions. We thank Jun Miao for providing the DiCre-dCas9 plasmids, Moritz Treeck for providing the DiCre-NF54 parasite line, Keith Joung for providing the high-fidelity Cas9, and Ashley Vaughan for providing the Linker3 plasmid and codon-optimized mNeonGreen plasmid.

This work was supported by funding from the National Institute of Allergy and Infectious Disease (F31 AI167494-01A1).

