## Supplemental Data for "A Conditional Cas9 System for Stage-Specific Gene Editing in *P. falciparum*"

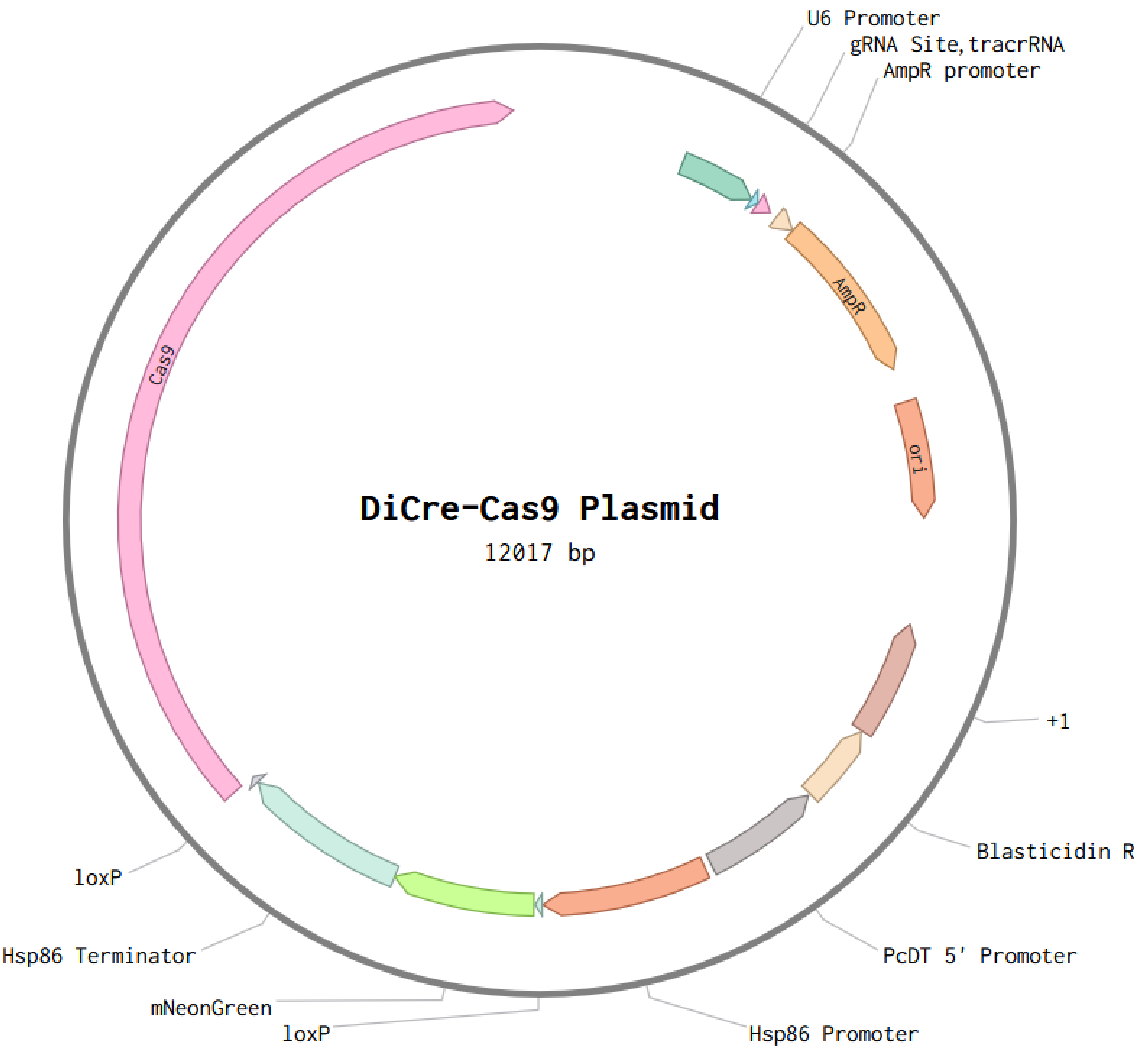


**Supplementary Figure 1.**

Plasmid map for the constructed DiCre-Cas9 plasmid. Contains: a U6 promoter followed by a gRNA cloning site and a tracrRNA, an Ampicillin resistance cassette, an origin of replication sequence, a Blasticidin resistance cassette driven by the PcDT promoter, an Hsp86 promoter, mNeonGreen and an Hsp86 terminator flanked by loxP sites, and a Cas9 following the loxP sites. The total plasmid size is 12,017 bp.

taaagggggcttaattataaaacagaaattattcttatcttacatgcacatataaaaaaatggattggtggtaaaccaaaaaaataaattcactatat
gttcttaaggaagcatataatgttttcctttttttttacatgcagatataaaaaggtagaagaacttacgagaagctctatattttacacatgcgatttgg
atatatatttttttttttgtaactttctatcatacttgtcataaattctgaattatcaaataactcaaatatatttcataatatcataatgctacaataataaa
aatatttttaatatatattaaaaaaaaaaattatatatatatttatatatatataatatacctatatacatacactatttttcattataattttttttttttttt
gtgtttttttatatatttttggaaatatgtaacgtataaaaaacaagacaaatataatatatactattaataaataagatatagttcttgttttaatattaat
taaaagaaaattttgtgaatatataaaaaaaaaaaaaaaaaaaaagaaaattataaataaatatatatattcgtgtaaaaataagtagaaaccac
gtatattataaattacaattcATGATGAGAAAATTAGCTATTTTATCTGTTTCTTCCTTTTTATTTGTTGAGGCCTTATTCCAG
GAATACCAGTGCTATGGAAGTTCGTCAAACACAAGGGTTCTAAATGAATTAAATTATGATAATGCAGGCACTAATT
TATATAATGAATTAGAAATGAATTATTATGGGAAACAGGAAAATTGGTATAGTCTTAAAAAAAATAGTAGATCACT
TGGAGAAAATGATGATGGAAATAACGAAGACAACGAGAAATTAAGGAAACCAAAACATAAAAAATTAAAGCAAC
CAGCGGATGGTAATCCTGATCCAAATGCAAACCCAAATGTAGATCCCAATGCCAACCCAAATGTAGATCCAAATGC
AAACCCAAATGTAGATCCAAATGCAAACCCAAATGCAAACCCAAATGCAAACCCAAATGCAAACCCAAATGCAAA
CCCAAATGCAAACCCAAATGCAAACCCAAATGCAAACCCAAATGCAAACCCAAATGCAAACCCAAATGCAAACCCA
AATGCAAACCCAAATGCAAACCCAAATGCAAACCCCAATGCAAATCCTAATGCAAACCCAAATGCAAACCCAAACG
TAGATCCTAATGCAAATCCAAATGCAAACCCAAACGCAAACCCCAATGCAAATCCTAATGCAAACCCCAATGCAAA
TCCTAATGCAAATCCTAATGCCAATCCAAATGCAAATCCAAATGCAAACCCAAACGCAAACCCCAATGCAAATCCTA
ATGCCAATCCAAATGCAAATCCAAATGCAAACCCAAATGCAAACCCAAATGCAAACCCCAATGCAAATCCTAATAA
AAACAATCAAGGTAATGGACAAGGTCACAATATGCCAAATGACCCAAACCGAAATGTAGATGAAAATGCTAATGC
CAACAGTGCTGTAAAAAATAATAATAACGAAGAACCAAGTGATAAGCACATAAAAGAATATTTAAACAAAATACA
AAATTCTCTTTCAACTGAATGGTCCCCATGTAGTGTAACTTGTGGAAATGGTATTCAAGTTAGAATAAAGCCTGGCT
CTGCTAATAAACCTAAAGACGAATTAGATTATGCAAATGATATTGAAAAAAAAATTTGTAAAATGGAAAAATGTTC
CAGTGTGTTTAATGTCGTAAATAGTTCAATAGGATTAATAATGGTATTATCCTTCTTGTTCCTTAATTAGataaagaac
acatcttagtttgagttgtacaatatttataaaaatatatactactttttttcttaattttcatttttctttatattttcctatttaatttattttttttgtgaatattt
aattacgtttgcgattaattgtagaaatatatatgtatatactatatttatagaatgtgttattctcaaaaacaacaacaaaaaaaaaaaaaaaaaaa
aaaaaaaagaaaaaaggattaaaagtaaaatagttataaatattttcaaaaatatttataacacaaaaaatacttcgaagttcatttaacatttttgtt
tatttatttatttatatatttcatttttacgtatttatattataaaatggtgtatcttaaaaatagtgaactatatatataaaatattaatttaaaaaaattat
aactttctttttattttctaaaataacttaaaaattatatgtttaagaaaggggtaaattataatatttgtataaatatataaacatagatatattaaataa
aataacaaatgtactatatttgtgcataagacgtatacgcttcatataatacatatatatatatatatatatatatatatattaattctat

**Supplementary Text 1.**
DNA sequence of CSP. CSP coding sequence is in capital letters, while non-coding sequence is in lowercase letters. The gRNA is highlighted in cyan. For the donor template, the 500bp homology arms are in light red, 200bp in red, and 80bp in dark red. Primers are highlighted in green.

taactatttttttttttttttttttttttttataaatttccccactcaaaATGAGGAATATCTTATTTATCCTCTCCTTTTTTTTGTGTGTTGTCT
ATGCACAAATTCCATATGATGTTCAAAGGATATATAAGGTTCTTATATTCTTATAAAAAAAAAAAAAAAACTACATAT
ATATATATATATATATATAATATTACTATATGTATCTTTCTATTATATTTTTCCGATAATGTTTTTATTTGTTGAATTGT
ATTTTTCTTTTATAGCCATCAAATCCTCAATCTGTTCATGCTCTTCCGGGAGACAAAGTTGCTTCAGGTTATAATCAA
AACAATCCATTCTTTCTTCCTTCATTTTGTAGTTTGTTAATTTTCTTTTTTCTTTGTTTCTTTTTAAGATTACAAATTTAA
GGATACTAAAATGGAAGTGGAAAAAGACAATAACAAGAATTCTTTGAAGGTATATTTTTCATTGTAGTAAATATGT
TAAAGCAACTCACTTTATAATGTTCCTATATATAATACGTGCAGTTAACATGTATATTTTTTATATGATGAATGTGGA
ATATATATATATATATAAATTTATTTATATAAATAAGTACATATATTATTTCCACCTTTTAGTGGTTAGCACAGAAGT
ATTACAAAGATGTCGATGAAACGAATAAAAAATATTTTGGAAGAATTAGATCAAGTAAAAATTTTAATCAAAATATA
TATATATATATATATATATATATATATGTGAATACACCATTTGCTTTTATTTATTATTTTTTTTTTTTAGCTTAGTGACAT
AATCGGGGAGAGAACATTATACTTCACTATATTTTAAataaatatactttttctataaaaactgtatgtgatataa

**Supplementary Text 2.**

DNA sequence of PF3D7_1316300. PF3D7_1316300 coding sequence is in capital letters, while non-coding sequence is in lowercase letters. The gRNA is highlighted in cyan. For the donor template, the homology arms are in red. Primers are highlighted in green.

tacataaacctgtatttaacctattcttttatgtctgaaattatacttgtttaatgatataatcgatttcctttttaacttatataaatattttatattattttac
ttattttattttatttattttatttttttttttttttgtaaaattgtttatATGATGCATCCCTTTAATTTTGTGCCTCAATTAGATAACA
AAATAAATGTTGTTTCAATGCAACCTTTCCAAATGTATGTTCCTAATAATAATGCTGTAATTCCTCAATCATTTTCCA
GTGATCATACAACACAACATTATAGTCAGCCTATATATTTTGAACCATTACCTCCTATATATGTGAAAAACCAATTAC
TACCTTCTCCCATTTTAGTACAGATGCCTACAACGGTAGTTGTTCAAAATGAATCACAACCAGCAATGGTCCTTAAT
CAACCACCCTCCAATATTGTTGTGAAAAATAATCCTCCAACTTCAGTTTTTGTTAAACAATCAAATCCTAATGTTGTT
GTTAAAAATGAAGCACCCCCAAATTATATACAAAATGTGGACACATGTCAAGAGACGATAGTAACAGATATGAAC
TGCCCAGTAATGACCAATATAAATAAAAATATTATGTAAtaagtgggaaataaatgactagacaattcaatatatgtcaaatggaaa
attgggattattattgaggcgttaatgtgatatacagaatgcaaatcgaaaactattgaaaaattatataattagaaataataaaaaa

**Supplementary Text 3.**

DNA sequence of PF3D7_1454900. PF3D7_1454900 coding sequence is in capital letters, while non-coding sequence is in lowercase letters. The gRNA is highlighted in cyan. For the donor template, the homology arms are in red. Primers are highlighted in green.

gcacatatttcatattaagtatataatatt[gRNA]gttttagagctagaaatagcaagttaaaataaggctagtccgttatcaacttgaaaaagtggc
accgagtcggtgcttttttt[Donor]gacgtcaggtggcacttttcgggga

**Supplementary Text 4.**

DNA sequence of a synthesized oligo for paired gRNA-donor templates. The 25-30bp homology arms for Gibson assembly cloning are in orange text. The 20bp variable gRNA is in cyan text. The 82bp tracrRNA is in purple text. The variable donor template is in blue text. Designed gRNAs should sit as close to the desired mutation as possible. Designed donor templates should be proximal to the gRNA cut site, and should disrupt the gRNA and/or NGG PAM site, either by deletion or by silent mutations.

| Primer | Dir. | Sequence | Len. | Tm | Type | Description |
|---|---|---|---|---|---|---|
| pSTW384 | FWD | TTTCTTCTTCAGGGTAGGCGGCCCTAATTTAATATAAAAAAAAATTCTTGCTTG | 54 | 60.0° | GA | Linker3 sequence to move U6, backbone, and donor template to DiCre plasmid |
| pSTW385 | REV | CATGGAACTCCTAGGCTGGGGTTATATAGCAAAAGAAAAGAAAGCGG | 47 | 61.0° | GA | Linker3 sequence to move U6, backbone, and donor template to DiCre plasmid |
| pSTW386 | FWD | CTTTCTTTTCTTTTGCTATATAACCCCAGCCTAGGAGTTCCATGGAGTTATC | 52 | 66.0° | GA | DiCre plasmid without backbone, to GA with Linker3 backbone, gRNA, donor |
| pSTW387 | REV | GAATTTTTTTTTATATTAAATTAGGGCCGCCTACCCTGAAGAAGAAAAGTCC | 52 | 65.0° | GA | DiCre plasmid without backbone, to GA with Linker3 backbone, gRNA, donor |
| pSTW301 | FWD | ATGGACTATAAGGACCACGACGGAGAC | 27 | 67.0° | GA | HFCas9 into DiCre Plasmid |
| pSTW302 | REV | AATAAGAAAAACGAACATTAAGCTGCCATATCCTTACTTTTTCTTTTTTGCCTGGCCGGC | 60 | 69.0° | GA | HFCas9 into DiCre Plasmid |
| pSTW356 | FWD | TGGGAATGGATGAATTGTATAAGTAATTATATAATATATTTATGTACTCACAATGGGGTC | 60 | 60.0° | GA | hsp 3' end to GA with mNeonGreen |
| pSTW357 | REV | GTTGTCCTCTTCTCCTTTAGATACCATGTCGACATAACTTCGTATAATGTATGCTATACG | 60 | 60.0° | GA | hsp 5' with loxp end to GA with mNeonGreen |
| pSTW362 | FWD | CGTATAGCATACATTATACGAAGTTATGTCGACATGGTATCTAAAGGAGAAGAGGACAA | 59 | 61.0° | GA | mNeonGreen to GA to loxp and hsp term |
| pSTW363 | REV | GACCCCATTGTGAGTACATAAATATATTATATAATTACTTATACAATTCATCCATTCC | 58 | 54.0° | GA | mNeonGreen to GA to loxp and hsp term |
| pSTW416 | FWD | GTTTTAGAGCTAGAAATAGCAAGTTAAAATAAG | 33 | 60.0° | GA | DiCre plasmid for Gibson gRNAs in |
| pSTW417 | REV | AATATTATATACTTAATATGAAATATGTGCATATAGG | 37 | 57.0° | GA | DiCre plasmid for Gibson gRNAs in |
| pSTW-seq018 | FWD | GATCGGCGACCAGTACGC | 18 | 62.0° | Seq | Cas9 pos. 900 |
| pSTW-seq056 | REV | TAGCCGGCGTAGCCGTTCTTG | 21 | 62.0° | Seq | Cas9 pos. 1130 |
| pSTW-seq028 | FWD | CCTATATGCACATATTTCATATTAAG | 26 | 48.0° | Seq | sgRNA |
| pSTW-seq029 | REV | CAAGTTGATAACGGACTAGCC | 21 | 53.0° | Seq | sgRNA |
| pSTW-seq011 | FWD | TCTCATGAGCGGATACATATTTGAATG | 27 | 49.0° | Seq | After U6, before Amp |
| pSTW-seq053 | FWD | GTACATTTGTTATTTTATTTAATATATCTATG | 32 | 54.0° | Seq | CSP outside |
| pSTW-seq054 | REV | GGGCTTAATTATAAAACAGAAATTATTC | 28 | 56.0° | Seq | CSP outside |
| pSTW-seq055 | FWD | GCCCTTAAGGCCTGCAGG | 18 | 59.0° | Seq | CSP donor inside |
| pSTW-seq058 | FWD | GATTTGCATTCTGTATATCACATTAACGC | 29 | 56.0° | Seq | PF3D7_1454900 outside |
| pSTW-seq059 | REV | CTGAAATTATACTTGTTTAATGATATAATCGATTTCC | 37 | 55.0° | Seq | PF3D7_1454900 outside |
| pSTW-seq066 | FWD | CGTTTCATCGACATCTTTGTAATACTTCT | 29 | 56.0° | Seq | PF3D7_1316300 outside |
| pSTW-seq067 | REV | GTGTGTTGTCTATGCACAAATTCC | 24 | 55.0° | Seq | PF3D7_1316300 outside |

**Supplementary Table 1.**

All primers used in this study. Dir. Denotes whether it is a forward or reverse primer. Sequence contains the complete primer sequences, including primer overhangs for Gibson assembly. Len. Denotes the primer length. Tm denotes the annealing temperature for the priming sequence. Type denotes how the primer is to be used, with GA meaning Gibson assembly cloning, and Seq meaning sequencing.
